# A macrophage subpopulation promotes airineme-mediated intercellular communication in a Matrix Metalloproteinase-9 dependent manner

**DOI:** 10.1101/2022.11.28.518037

**Authors:** Raquel Lynn Bowman, Daoqin Wang, Dae Seok Eom

## Abstract

Tissue-resident macrophages are highly heterogenous and perform various dedicated functions depending on their locations. In particular, skin resident macrophages have intriguing roles in long-distance intercellular signaling by mediating cellular protrusions called ‘airinemes’ in zebrafish. During pigment pattern formation, macrophages relay signaling molecules containing ‘airineme vesicles’ from one pigment cell to another. Without macrophages, airineme-mediated signaling is abolished, disrupting pigment pattern formation. It remains unknown, however, if the same macrophage population controls both these signaling roles and typical immune functions or if a separate macrophage subpopulation functions in intercellular communication. In this study, with high-resolution confocal live-imaging and cell type-specific genetic ablation approaches *in vivo*, we have identified a macrophage subpopulation responsible for airineme-mediated signaling. These cells appear distinct from conventional skin resident macrophages by their amoeboid morphology and faster/expansive migratory behaviors. Instead, we show that they resemble ectoderm-derived macrophages termed metaphocytes. Metaphocyte ablation dramatically reduces airineme extension and signaling. In addition, these amoeboid/metaphocytes require high levels of MMP9 expression for their migration and airineme-mediated signaling. These results reveal a novel macrophage subpopulation with specialized functions in airineme-mediated signaling, which may play roles in many other aspects of intercellular communication.

## Introduction

Intercellular signaling in multicellular organisms is essential for development and homeostasis. Uncontrolled regulation of such signaling causes various human diseases, and altered intercellular communication is a hallmark of aging (Lopez-Otin et al., 2013). Intercellular communication mechanisms include endocrine, paracrine, juxtacrine, and neuronal signaling, with diffusion-based local signaling thought to be a dominant dissemination mechanism in paracrine signaling (Lander et al., 2002; Muller and Schier, 2011; Wolpert, 1973, 2009). In recent years, growing evidence has suggested that cells can communicate with each other via long, thin cellular protrusions extended by signal sending or receiving cells (Eom et al., 2015; Ljubojevic et al., 2021; Roy et al., 2014; Stanganello and Scholpp, 2016). These can be classified based on their cytoskeletal composition, mode of signal delivery, morphology, and more. The growing list of such protrusions that have been identified include cytonemes, tunneling nanotubes, airinemes, intercellular bridges, and migrasomes (Caviglia and Ober, 2018; Daly et al., 2022; Gonzalez-Mendez et al., 2019; Jiang et al., 2019). They are found in a variety of cell types and species, such as fruit fly, sea urchin, zebrafish, and cultured mammalian cells, and their signaling roles have been functionally validated (Eom, 2020). These signal-carrying protrusions are orders of magnitude longer than typical filopodia and can extend or retract in a highly dynamic fashion. They directly contact target cells and deliver the major morphogens in many paracrine contexts. In addition to diffusible signals, membrane ligand and receptor-mediated intercellular communication, such as Delta-Notch signaling, can also be mediated over long distances via such signaling cellular protrusions (Cohen et al., 2010; De Joussineau et al., 2003; Eom et al., 2015; Huang and Kornberg, 2015).

Previously we identified a class of signaling cellular protrusion called ‘airinemes’ critical for interactions between zebrafish pigment cells. They are unique in having large vesicle-like structures at their tips (Eom et al., 2015). They are also highly curved, and mathematical models suggest that such complex shapes are optimized to search for target cells (Park et al., 2022). Airinemes preferentially extend from undifferentiated yellow pigment cells, xanthoblasts, and their vesicles contain Delta C ligand, which they deliver to melanophores. Notch activation in target melanophores induces them to migrate from prospective interstripe to stripes, establishing stripe pigment formation. When airineme extensions are inhibited target melanophores are retained in the interstripe, which leads to failure of proper stripe consolidation. Intriguingly, we have also found that skin-resident macrophages play an essential role in airineme-mediated signaling. Skin-resident macrophages recognize the bulged membrane blebs from the surface of airineme-producing xanthoblasts. These blebs, precursors of airineme vesicles, are grabbed and extracted by macrophages, and as they migrate, filaments trail behind. Macrophages drag the vesicles along as they migrate through the tissue, deposit them onto the surface of target melanophores, and continue to wander. Thus, the highly curved airineme trajectories observed reflect complex migratory paths of skin-resident macrophages. Deposited vesicles on target melanophores then remain for over an hour or longer even after filament retraction, an interval presumed to be sufficient for activation of Notch signaling. Afterwards, other macrophages phagocytose the deposited airineme vesicles. Thus, macrophages seem to initiate and terminate airineme-mediated long-range Notch-Delta signaling (Eom, 2020; Eom et al., 2015; Eom and Parichy, 2017).

Macrophages are well described immune cells that phagocytose foreign pathogens and apoptotic bodies, critical for self-defense and maintaining homeostasis (Guilliams et al., 2020; Herbomel and Levraud, 2005; Herbomel et al., 1999; Varol et al., 2015). Tissue-Resident Macrophages (TRMs) are highly heterogeneous immune populations found in different local niches with unique tissue-specific functions. Unlike conventional macrophages derived from hematopoietic stem cells, TRMs originate in various tissues as shown by fate mapping studies in mice (Ginhoux and Guilliams, 2016; Perdiguero et al., 2015; Schulz et al., 2012). In zebrafish, ectoderm-derived skin resident macrophages are called ‘metaphocytes.’ Endoderm-derived metaphocytes are also found in the gills and intestine of zebrafish. While metaphocytes express phagocytic genes, they seem not professional phagocytes. Instead, they are specialized for capturing external soluble antigens and transferring them to other immune cells (Lin et al., 2020; Lin et al., 2019).

Prior studies have suggested that occasionally the specialized functions of TRMs are not directly involved in immunity (Bleriot et al., 2020). During development, they play essential roles in tissue remodeling (Bohaud et al., 2021; Eom and Parichy, 2017). Microglia, macrophages in the postnatal brain in mice, for example, are critical for synaptic pruning (Paolicelli et al., 2011). Other roles for TRMs include blood vessel pruning and regression and fin regeneration in zebrafish (Korn and Augustin, 2015; Petrie et al., 2014). TRMs in skin are essential in airineme-mediated intercellular signaling during postembryonic tissue remodeling (Eom, 2020; Eom and Parichy, 2017).

Both typical and atypical functions of macrophages depend on their ability to migrate. Macrophages can infiltrate most tissues and migrate between cells *in vivo* by degrading extracellular matrix (ECM) with extracellular proteinases such as matrix metalloproteinases (MMPs), cathepsins, and urokinase-type plasminogen activator (Verollet et al., 2011). A few studies have shown that MMP9 activity is critical in macrophage migration in mice and zebrafish (Gong et al., 2008; Travnickova et al., 2021).

Here we use high-resolution live imaging and genetic cell ablation to show that a skin-resident macrophage subpopulation resembling metaphocytes is essential in airineme-mediated intercellular communication between pigment cells in zebrafish. Consistent with this hypothesis, metaphocyte-specific ablation abrogates airineme extension and subsequent signaling. Lastly, we show that the ability of these macrophage subpopulations to mediate airineme signaling depends on MMP9 activity. Our study shows macrophage heterogeneity and the diversity of cellular functions performed by tissue-resident macrophages in cellular protrusion-mediated intercellular signaling.

## Results

### Two distinct macrophage subpopulations in zebrafish skin

Skin-resident macrophages play essential roles in airineme-mediated long-range intercellular communication (Eom, 2020; Eom and Parichy, 2017). Here we asked if the same macrophage population performs both its typical innate immune functions and the airineme-mediated signaling roles or if there are other subpopulations of macrophages specialized for airineme signaling. To look for distinct macrophage subpopulations in the zebrafish skin, we first examined *mpeg1+* macrophages using a transgenic reporter line, *Tg(mpeg1:tdTomato)*, at the metamorphic stages at which macrophage/airineme-mediated signaling most frequently occur (Eom et al., 2015). High-resolution, confocal time-lapsed imaging revealed two morphologically distinct *mpeg1+* macrophages. The first type includes cells that are relatively large, dendritic and express a higher level of *mpeg1*, while the second type is relatively smaller, amoeboid in shape and expresses less *mpeg1* (Fig. 1A, Supple Video 1). These two subpopulations also appeared to traverse different depths of the skin, which led us to ask how deep each population traveled, given that airineme-producing xanthoblasts are restricted to the hypodermis. For this, we measured the distance between each macrophage subpopulation and xanthoblasts. While both subpopulations were spread throughout the skin, there were significantly more amoeboid cells located basally in the hypodermis than dendritic macrophages (Fig. 1A), which were located apically near xanthoblasts (Fig. 1A, 1B, arrow). Based on their morphology and relative locations, amoeboid and dendritic macrophages appear to be two distinct subpopulations which led us to ask if these two subpopulations also behave differently. Studying their migration patterns revealed that amoeboid macrophages migrate significantly faster than dendritic macrophages (Fig. 1C). Strikingly, the migration speed of the amoeboid macrophages (4.7 μm/min) matched with the speed of airineme extension (4.7 μm/min) indicating that amoeboid macrophages were more likely to interact with airineme vesicles than dendritic macrophages at these metamorphic stages (Eom et al., 2015).

**Figure 1.**
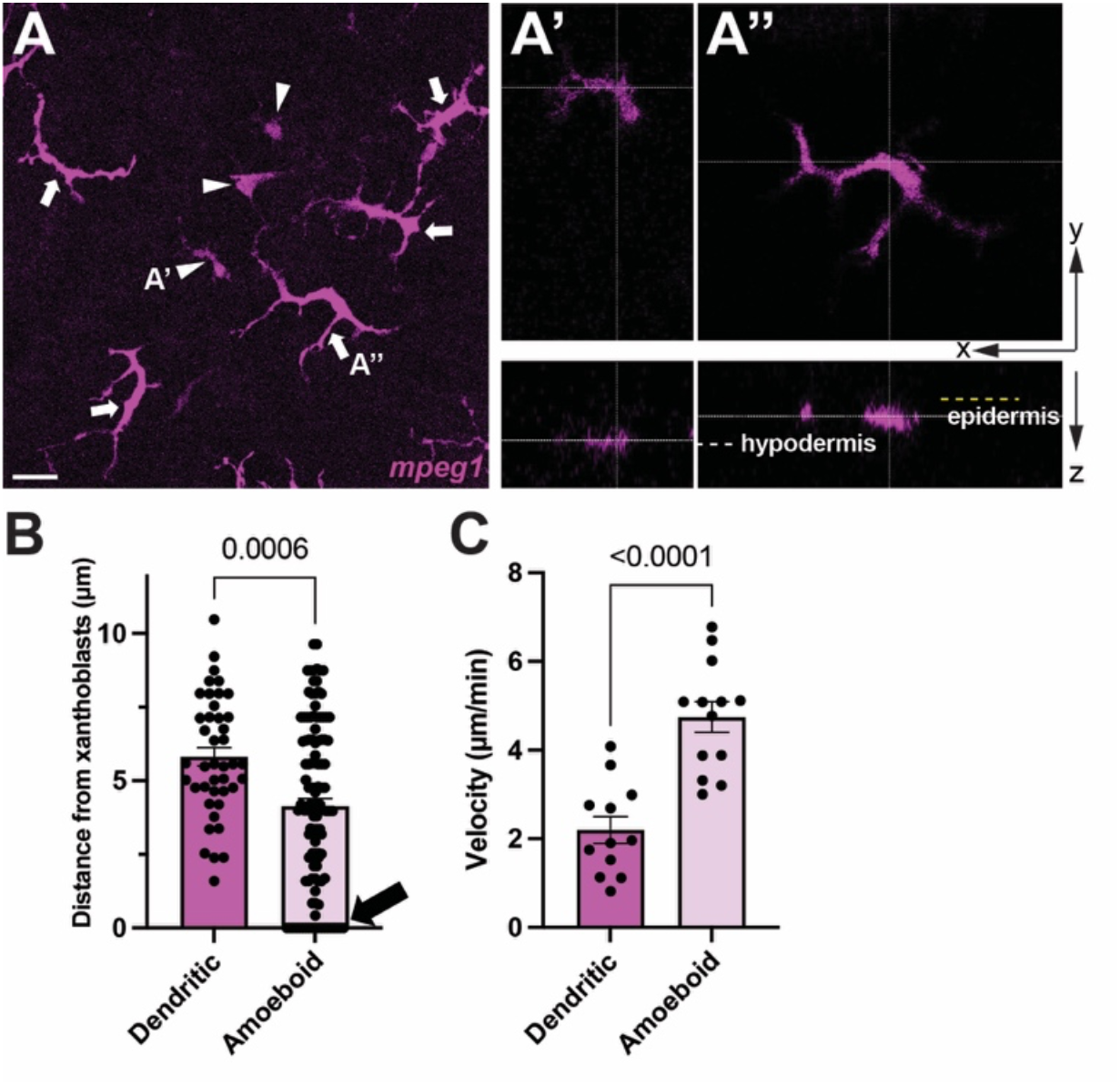
Two distinct macrophage subpopulations in zebrafish skin. (A) *mpeg1+* amoeboid (arrowheads) and dendritic (arrows) macrophages shown in maximum projection image. A’, amoeboid macrophages at hypodermis, A’’, dendritic macrophages at epidermis. Yellow dotted line denotes apical margin of epidermis. White dotted line denotes hypodermal layer. (B) Amoeboid macrophages are localized more basally than dendritic and reach hypodermis (arrow); n=63, 122 cells. (C) Migration speed of dendritic and amoeboid macrophages; n=12, 13 cells. Statistical significances were accessed by student t-test. Scale bars: 20μm.

### Amoeboid *mpeg1+* macrophages interact with airineme vesicles

Indeed, *ex-vivo* high-resolution live confocal imaging revealed that amoeboid macrophages almost exclusively interacted with airineme vesicles and pull them (∼96.8%) as compared to dendritic macrophages (∼3.125%) (Fig. 2A, B, Supple Video 2, arrowhead). Overall, our observations suggest that among two distinct macrophage subpopulations in zebrafish skin, the amoeboid macrophage subpopulation preferentially interacts with airinemes.

**Figure 2.**
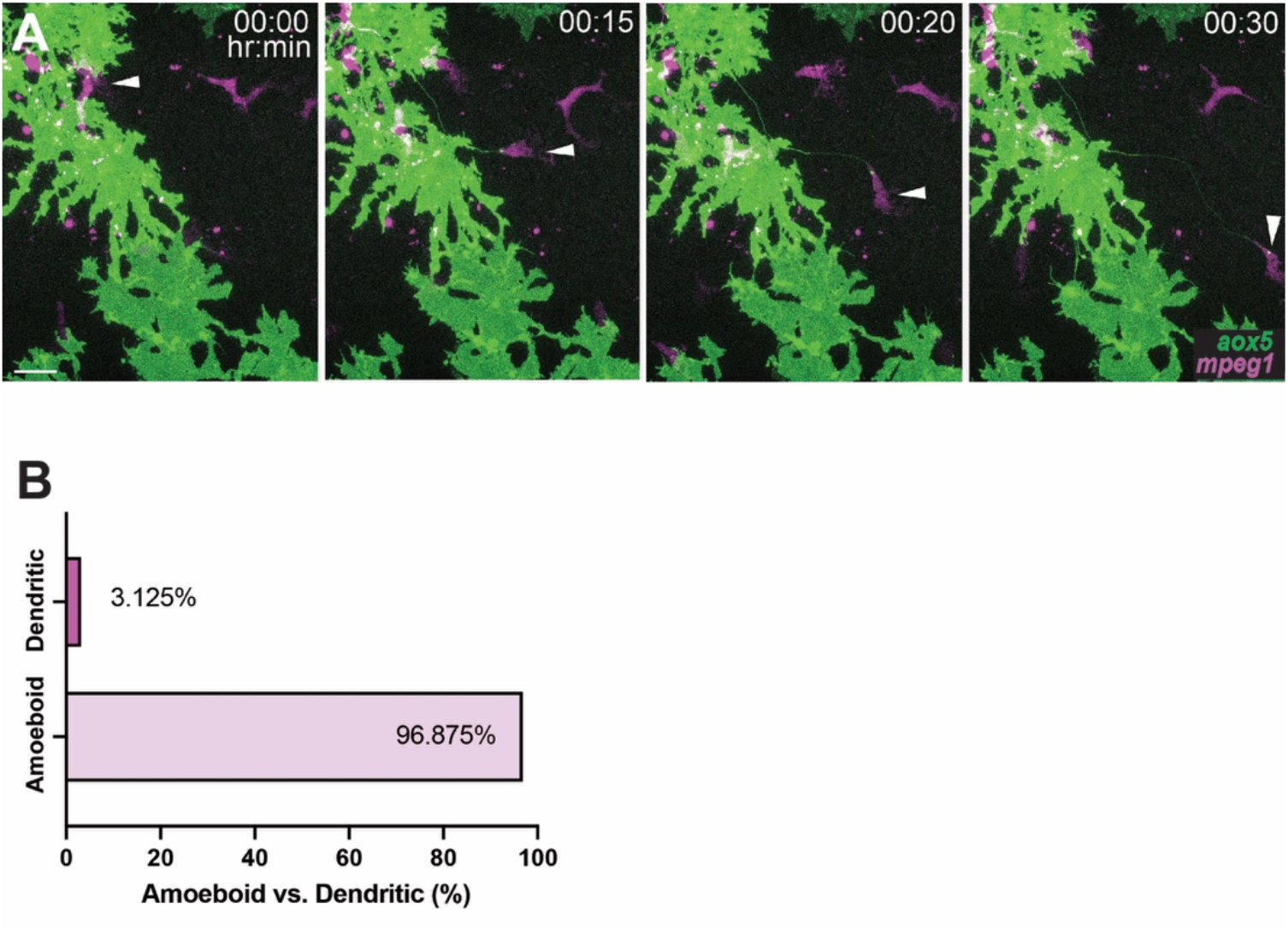
Amoeboid *mpeg1+* macrophages interact with airineme vesicles. (A) Airinemes are pulled by amoeboid macrophages (arrowhead). Still images from time-lapse movie. (B) Percentages of airineme-pulling macrophage subpopulations. Scale bars: 20μm.

As mentioned, airinemes extend most frequently during metamorphic stages (SSL 7.5) (Eom et al., 2015). Thus, we asked if the amoeboid subset of macrophages proportionally expands at these stages. However, both overall skin-resident macrophage (*mpeg1+*) numbers and the amoeboid subpopulation increased in similar proportions, with the ameboid population maintained at 24-29% of total macrophages during SSL5.5∼8.5 (Supple Fig. 1A).

### Airineme-pulling amoeboid macrophages overlap with metaphocytes

Traditionally, macrophages have been classified largely into two subpopulations based on their polarization (Davies et al., 2013; Guilliams et al., 2020; Rostam et al., 2017). Classically activated M1 macrophages are round, while alternatively activated M2 macrophages are more elongated (Varol et al., 2015; Vereyken et al., 2011). To determine if airineme-associated amoeboid macrophages are more M1- or M2-like we examined the M1 marker, *TNFα*. However, neither amoeboid nor dendritic macrophages expressed this marker (Supple Fig. 2A). This result is consistent with our previous finding that airineme extension is independent of macrophage activation (Eom et al., 2015).

Ectoderm- or endoderm-derived macrophages termed ‘metaphocytes’ have been recently identified in zebrafish (Kuil et al., 2020; Lin et al., 2020; Lin et al., 2019). Metaphocytes are transcriptionally similar to conventional tissue-resident macrophages in the epidermis (Lin et al., 2019) and comparable to conventional macrophages, which originate during hematopoiesis (Davies et al., 2013). Interestingly, it has been suggested that metaphocytes function by delivering antigens rather than responding to immune-related events. Moreover, their morphology resembles that of the airineme-pulling amoeboid macrophage subpopulation (Fig. 1A-A’’, Fig. 2A) (Kuil et al., 2020; Lin et al., 2020; Lin et al., 2019). Thus, we hypothesized that metaphocytes promote airineme-mediated intercellular signaling. We first tested if amoeboid macrophages express metaphocyte markers. Metaphocyte-specific reporter lines in which EGFP expression is driven by the promoter of either *claudin-h (cldnh)* or *granulin2 (grn2)* were crossed with the macrophage reporter, *Tg(mpeg1:tdTomato)* (Lin et al., 2020; Lin et al., 2019). *Tg(cldnh:EGFP; mpeg1:tdTomato)* and *Tg(grn2:EGFP; mpeg1:tdTomato)* both label amoeboid *mpeg1+* macrophages, while dendritic populations do not express either metaphocyte marker (Fig. 3A, arrowheads, Supple Fig. 2B). In addition, the number of amoeboid macrophages and metaphocytes *(grn2+/mpeg1+)* at each developmental stage were similar, suggesting that the airineme interacting amoeboid macrophage subpopulation largely overlaps with metaphocytes in zebrafish skin (Supple Fig. 1B).

**Figure 3.**
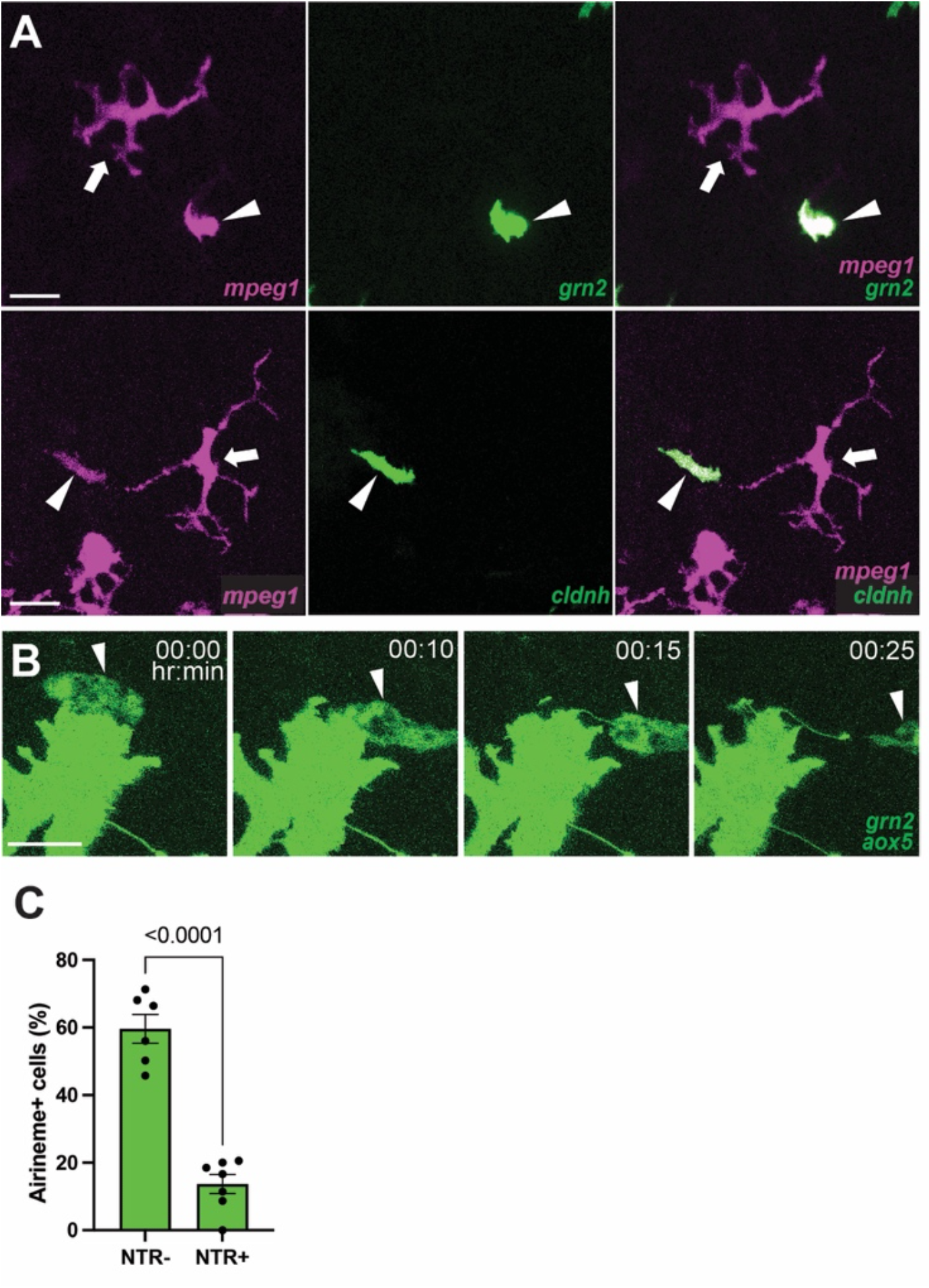
Airineme pulling amoeboid macrophages overlap with ectoderm-derived metaphocytes. (A) Coexpression of metaphocyte markers, *grn2* and *cldnh*, in amoeboid macrophage subpopulation (*mpeg1+*, arrowheads). Note that dendritic (*mpeg1+*, arrows) macrophages do not express metaphocyte markers. (B) Xanthoblasts of metaphocyte depleted fish (NTR+) were less likely than controls (NTR-) to extend airinemes (n=6, 7 time-lapse positions, 3 trunks each). Statistical significances were accessed by student t-test. Scale bars: 20μm (A), 10μm (B).

We confirmed that metaphocytes associate with airinemes with *ex vivo* live imaging (Fig. 3B, Supple Video 3, arrowhead). We hypothesized that if metaphocytes are an airineme-pulling macrophage subpopulation, we expect a significant inhibition of airineme extension in the absence of metaphocytes. To test this idea, we specifically ablated metaphocytes by utilizing a transgenic line, *Tg(grn2:EGFP-v2a-nfsB)* that expresses the enzyme nitroreductase (NTR, *nfsB*) in metaphocytes. NTR converts the innocuous prodrug metronidazole (Mtz) into a toxic metabolite that kills cells (Curado et al., 2008; Pisharath and Parsons, 2009). In metaphocyte-specific depleting conditions (see Methods), we observed that airineme extension was significantly reduced (Fig. 3C). Taken together, we conclude that the amoeboid macrophage subpopulation overlaps with metaphocytes, and they are responsible for airineme-mediated signaling.

### Metaphocyte-mediated airineme extension is responsible for pigment pattern development

Airineme-mediated signaling is critical for pigment pattern formation. During metamorphosis, signal molecule-bearing airineme vesicles are relayed to target melanophores by macrophages. The target cells receive signals from airineme vesicles and are directed to migrate out of the interstripe and coalesce into stripes (Eom, 2020; Eom et al., 2015; Eom and Parichy, 2017). Thus, we asked if metaphocytes are responsible for pigment pattern formation. We ablated metaphocytes continuously during pigment pattern development. Repeated metaphocyte ablation exhibited melanophore retention in interstripe, while the controls had few to no melanophores at SSL 12 (see Methods). Total numbers of melanophores were not altered in either the controls or the experimental, suggesting target melanophore migration into stripes was hindered in the absence of metaphocytes (Fig. 4A, B). The disorganized pigment patterns observed resembled the phenotypes found when airinemes were inhibited, or all *mpeg1+* macrophages in the skin were depleted (Eom et al., 2015; Eom and Parichy, 2017). Our data suggest that metaphocytes are responsible for pigment pattern formation via airineme-mediated signaling.

**Figure 4.**
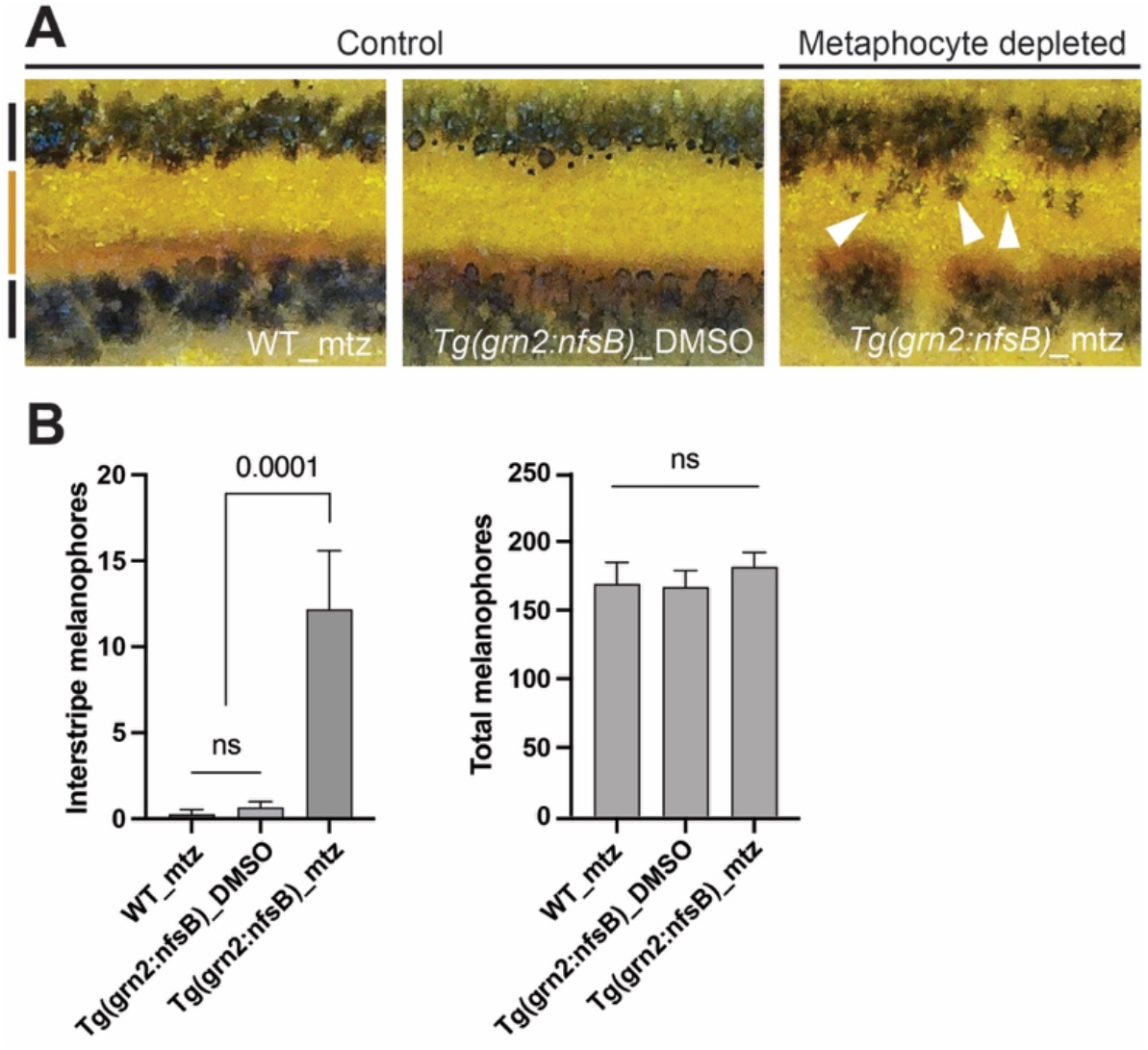
Metaphocyte-mediated airineme extension is responsible for melanophore pattern development. (A) In two controls, melanophores reside in the stripe region (denoted by black bars at far left) and border the interstripe (orange bar). Metaphocyte depletion results in melanophores retention in the interstripe (arrowheads). (B) Numbers of melanophores in the interstripe. Metaphocyte depleted fish (*Tg(grn2:NTR)_*mtz) had significantly more melanophores in the interstripe than controls (WT_mtz and *Tg(grn2:NTR)_*DMSO) and displayed disorganized pigment pattern, although total melanophore numbers did not differ; data are means ± SE. At stage 12 standardized standard length (SSL) (n=4, 15, 11 trunks). Statistical significances were accessed by student t-test.

### Metaphocyte migration and airineme extension are MMP9 dependent

We asked what underlying molecular mechanism differentiates amoeboid/metaphocytes from the dendritic macrophage subpopulation. Matrix Metalloproteinases (MMPs) are a critical component of macrophage migration. Among many MMPs, MMP9 has been suggested to be required for macrophage migration and phagocytosis (Gong et al., 2008; Travnickova et al., 2021). Therefore, we hypothesized that the differential MMP9 requirement governs migratory behavior between amoeboid/metaphocytes and dendritic macrophages in zebrafish skin. First, we tested if mmp9 is differentially expressed in these two macrophage subpopulations. We generated a double transgenic line, *Tg(mmp9:EGFP; mpeg1:tdTomato)*, that labels both macrophage subpopulations and mmp9-positive cells (Ando et al., 2017). We observed that amoeboid/metaphocytes exhibited strong mmp9 expression, but the dendritic population showed no detectable reporter expression (Fig. 5A, Supple Fig. 2B, 2C). Consistently, only amoeboid/metaphocyte migration speed was significantly reduced in the presence of MMP9 inhibitor. However, dendritic cell migration was unchanged, suggesting that MMP9 is indispensable for amoeboid/metaphocyte migration (Fig. 5B, Supple Fig. 3). Consistent with airinemes interacting specifically with amoeboid/metaphocytes, we observed similar reductions in airineme extension speed in the presence of an MMP9 inhibitor (Fig. 5C). Surprisingly, we discovered that airineme extension frequency was significantly compromised upon treatment with this MMP9 inhibitor (Fig. 5D). We reasoned this might be due to the drug preventing metaphocyte infiltration into the hypodermis. Indeed, we observed that the localization of amoeboid/metaphocytes along the apical-basal axis was shifted apically in the presence of MMP9 inhibitor compared with controls. Thus, limited access of amoeboid/metaphocytes to the hypodermal layer could cause the significant reductions in airineme extension frequency seen in MMP9 inhibitor-treated animals (Fig. 5E). Lastly, we observed that the airineme extension was significantly reduced when we ablated *mmp9+* cells with *Tg(mmp9:nfsB-EGFP)* (Ando et al., 2017) (Fig. 5F). Together, these data suggest that amoeboid/metaphocyte-mediated airineme signaling depends on MMP9 activity.

**Figure 5.**
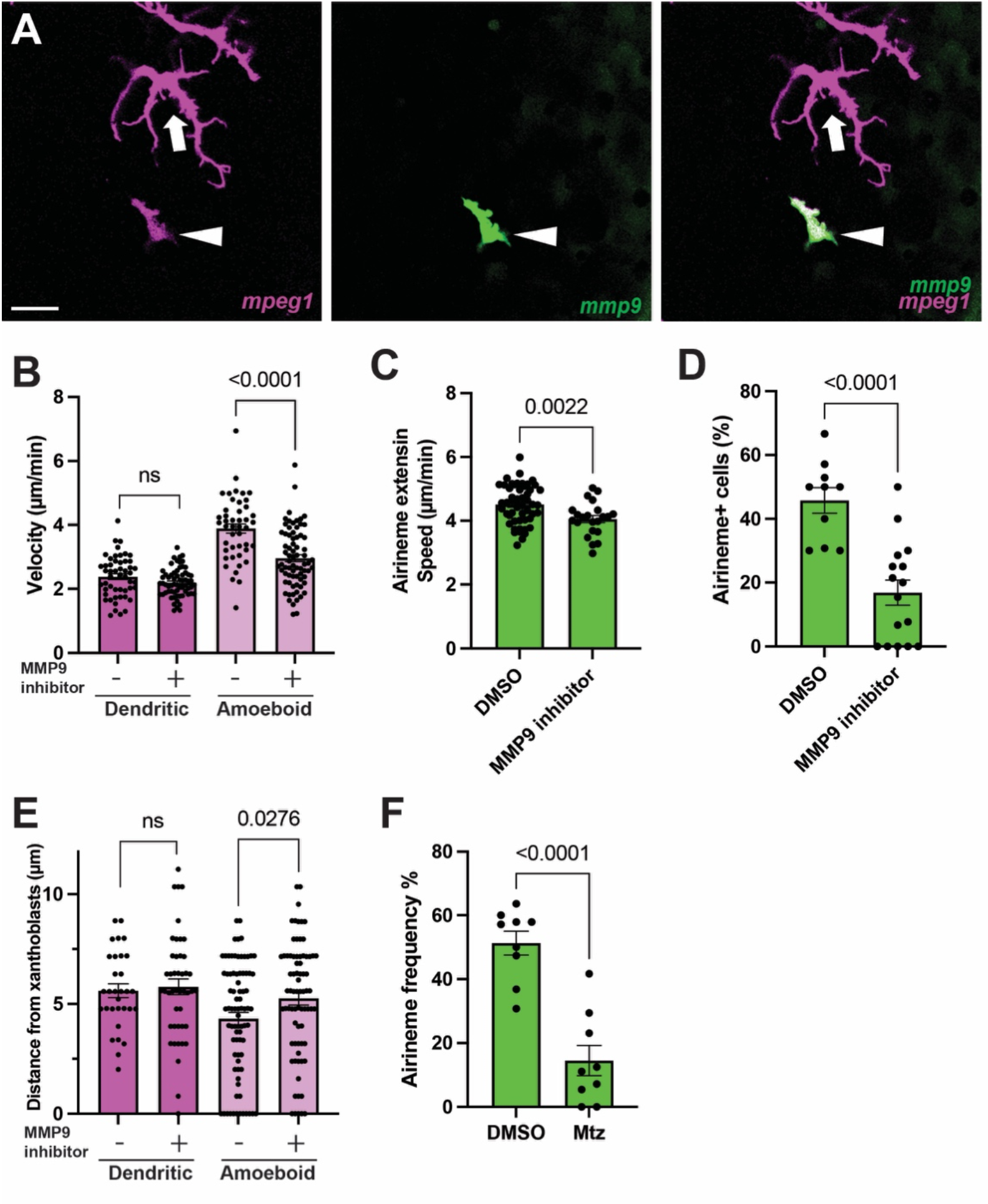
Metaphocyte migration and airineme extension are MMP9 dependent. (A) Differential expression of mmp9 in amoeboid (arrowhead) and dendritic macrophage subpopulation (arrow). (B) MMP9 inhibitor treatment significantly reduced the migration speed of amoeboid macrophage subpopulation but not dendritic macrophages (n=51, 64, 45, 68, 4 trunks each). (C) Airineme extension speed reduction in the presence of MMP9 inhibitor (n=53, 23, 3 trunks each). (D) Xanthoblasts in MMP9 inhibitor treated fish had fewer airinemes than controls (n=10, 16 time-lapse positions, 4, 5 trunks). (E) Localization of amoeboid macrophages were shifted more apically upon MMP9 inhibitor treatment but no effects on dendritic macrophage subpopulation (n=31, 45, 80, 75 cells, 3, 3, 4, 5 trunks). (F) Xanthoblasts in mmp9+ cell depleted fish had fewer airinemes than controls (n=9,9 time-lapse positions, 3 trunks each). Scale Bars, 20μm (A). Statistical significances were accessed by student t-test.

## Discussion

Several features of airinemes distinguish them from other signaling cellular protrusions, such as cytonemes and tunneling nanotubes. One of the most striking differences is that airinemes require macrophages as ‘delivery vehicles’ for relaying airineme vesicles to the target melanophores intermingled with non-target cells (Daly et al., 2022; Eom, 2020). Airinemes may need these cells to help them extend through the densely packed tissue environment, especially airinemes with large vesicles at their tips (Hirata et al., 2003).

Our study suggests that amoeboid macrophages/metaphocytes in metamorphic zebrafish skin specifically perform such a delivery role in airineme signaling. We have shown that the behaviors of this macrophage subpopulation correlate with airineme-mediated signaling since they migrate faster and can infiltrate deeper into hypodermis than the dendritic subpopulation (Fig. 6). Our study also suggests that MMP9-dependent migration of metaphocytes is critical for airineme-mediated intercellular communication. However, dendritic macrophages do not seem to express MMP9, and their migration was not affected by MMP9 inhibitor treatment (Fig. 5). There are 26 MMP-orthologs in zebrafish and 24 different MMP-genes in humans. Thus, it will be interesting to study whether combinatorial MMP expression control or fine-tune macrophages’ migratory behaviors in different subpopulations or disease contexts. This idea is supported by studies showing that MMP-mediated macrophage infiltration is associated with various human pathophysiology (Lagente et al., 2009; Nighot et al., 2021; Pedersen et al., 2015).

**Figure 6.**
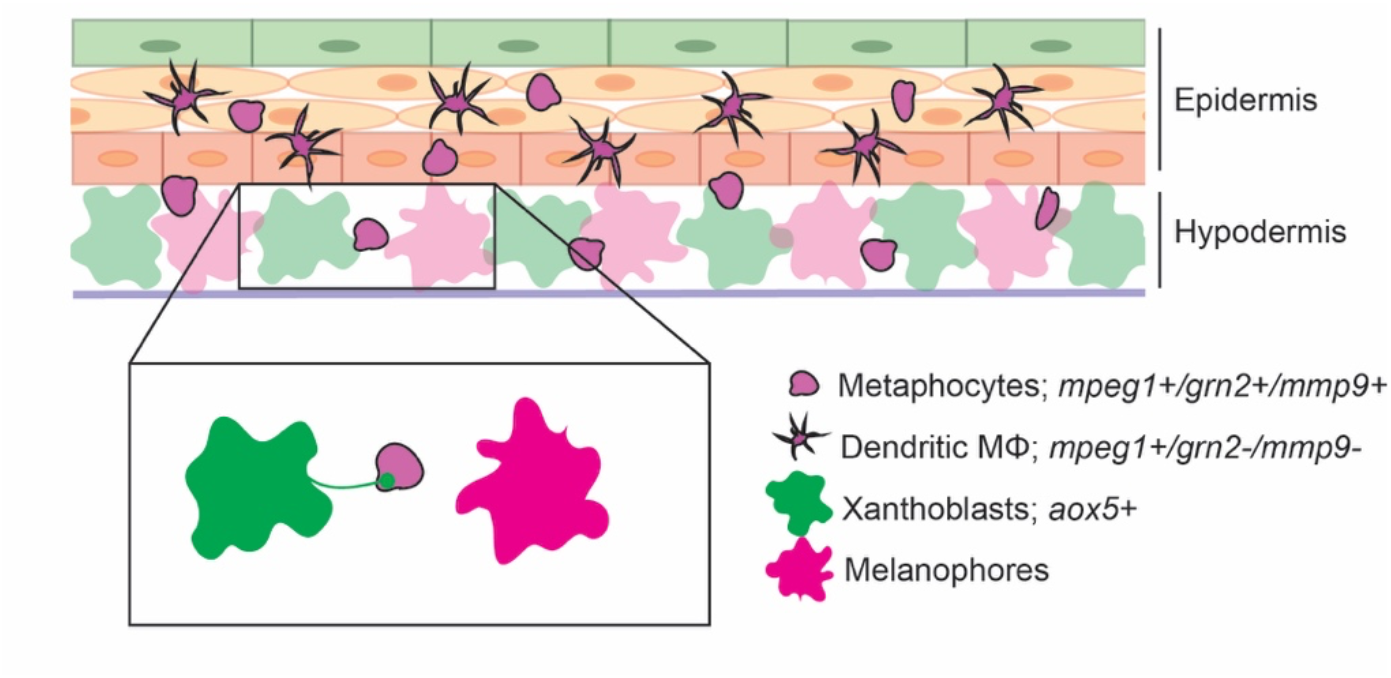
Amoeboid/metaphocytes-mediated airineme signaling. Unlike the dendritic macrophage subpopulation (*mpeg1+/grn2-/mmp9-*), amoeboid macrophages/metaphocytes are found in both epidermis and hypodermis at the metamorphic zebrafish skin. Amoeboid macrophage subpopulation/metaphocytes pull airinemes at hypodermis via MMP9 dependent manner.

Our results are consistent with previous reports showing that amoeboid/metaphocytes are specialized for antigen delivery but less reactive to immunogenic processes (Lin et al., 2019). Metaphocytes in metamorphic zebrafish skin is also specialized for airineme vesicle delivery. However, from our previous studies, we observed that metaphocytes engulf and eliminate the airineme vesicles when they are docked on non-target cells suggesting they are still phagocytes. However, airineme vesicles are not phagocytosed while airinemes extend and when they contact target cells (Eom et al., 2015; Eom and Parichy, 2017). Thus, it appears that the target recognition mechanism is tightly linked to determining metaphocytes to phagocytose the airineme vesicles. We speculate that these amoeboid/metaphocytes control target recognition in airineme-mediated intercellular communication. One possibility is that metaphocytes scan the surfaces of cells along their migration paths and deposit airineme vesicles onto specific target melanophores. Thus, cell membrane receptor/ligand interactions between metaphocytes and target cells might provide a cue for target cell specificity, and this might prevent airineme vesicle phagocytosis. Alternatively, it is possible that interactions between airineme vesicles and target cell membranes underlie this target specificity and metaphocytes simply act as delivery vehicles. However, the molecular mechanisms behind metaphocytes’ target specific behaviors remain to be elucidated.

In this study, we identified two distinct macrophage subpopulations in metamorphic zebrafish skin, consistent with previous descriptions of metaphocytes in adult zebrafish (Lin et al., 2020; Lin et al., 2019). In contrast, however, the localization of metaphocytes in adults is near the peridermal epidermis and conventional dendritic macrophages stay relatively more basally, while we find that our amoeboid, metaphocyte-like cells lie more basally than dendritic macrophages (Fig. 1 & 6). This discrepancy may simply reflect stage-dependent differences in these cell types. In our study, we have shown that airinemes are most frequently observed during metamorphosis where robust tissue remodeling, including pigment pattern formation, occurs, but their extension frequency diminishes as fish develop (Eom et al., 2015). Thus, it is conceivable that metaphocytes repurpose their function to relay antigens in adult zebrafish skin.

It is important to note that in rare cases we observed amoeboid macrophages which did not express metaphocyte markers, *grn2 or cldnh*. We also observed that *grn2+* metaphocytes displayed a varying degree of reporter expression in the transgenic lines. This may reflect their maturation status or suggest the existence of additional macrophage subpopulations. However, these non-metaphocyte amoeboid macrophages make up a much smaller portion than most amoeboid/grn2+ metaphocytes in metamorphic zebrafish skin.

Taken all together, our study revealed that airineme-vesicles interact specifically with an amoeboid macrophage subpopulation, most likely ectoderm-derived macrophages called metaphocytes. Moreover, metaphocytes are responsible for airineme-mediated signaling via MMP9 activity-dependent fashion (Fig. 6). Our study suggests that macrophage involvement in airineme-mediated signaling is not an experimental side effect but highly regulatory and that airinemes achieve intercellular signaling by utilizing tissue-resident macrophage heterogeneity and versatility. Our study also demonstrates the complex and cooperative nature of the cellular protrusion-based intercellular signaling between pigment cells in zebrafish skin. Requirements for macrophages in other cellular protrusion-mediated signaling modalities are unknown, which is an interesting avenue for future studies.

## Materials and methods

### Staging, Rearing, and Stocks

Fish were maintained at 28.5 C, 16:8 L:D. Zebrafish were wild-type AB^*wp*^ or its derivative WT (ABb), as well as, *Tg(cldnh:EGFP), Tg(grn2:EGFP;mpeg1*.*1:LoxP-DSRedx-LoxP-EGFP), Tg(grn2:EGFP-v2a-nfsB)*^*hkz034*^, provided by Z. Wen (Lin et al., 2020; Lin et al., 2019), *Tg(tyrp1b:palmmcherry)*^*wp*.*rt11*^ (Eom et al., 2015), *Tg(mpeg1:Brainbow)*^*w201*^, which expresses tdTomato in the absence of Cre-mediated recombination (Pagan et al., 2015); *Tg(mmp9:EGFP)*^*tyt206*^, *Tg(mmp9:EGFP-v2a-nfsB)*^*tyt207*^, provided by A. Kawakami (Ando et al., 2017). Experiments were performed prior to the development of secondary sexual characteristics, so the number of females and males used in the study could not be determined, however, all stocks used generated approximately balanced sex ratios, so the experiments likely sampled similar numbers of females and males.

### Transgenesis and transgenic line production

*Tg(cldnh:eYFP-v2a-nfsB):* To ablate cldnh labeled cells in a cell specific manner, we generated *cldnh:eYFP-v2a-nfsB* construct from the *cldnh:EGFP* construct provided by Z. Wen (Lin et al., 2019). Briefly, a Gateway middle entry fragment *(pME:eYFP-v2a-nfsB)* was generated using the Gibson assembly (Gibson et al., 2009). Later, *pME:eYFP-v2a-nfsB* was combined with *cldnh* promoter backbone in an In-Fusion reaction to generate *pBLK-cldnh:eYFP-v2a-nfsB*. The resulting construct also included a Tol2 sequence which allowed for utilization of the Tol2 transposon system to randomly incorporate our plasmid of interest into WT(ABb) fish.

*cldnh:mcherry* construct: The same *cldnh:EGFP* construct was subcloned to generate *cldnh:mCherry* to label *cldnh* positive cells other than green fluorescent protein using Gibson assembly (Gibson et al., 2009).

### Drug treatments

Metaphocyte Long-Term Ablation: Fish were treated with drugs during dark cycles and then washed and changed into system water without drugs at light cycles. *Tg(grn2:EGFP-v2a-nfsB)* fish were split into two groups: DMSO control and metronidazole (Mtz)-treated experimental and WT(ABb) were used as an additional control group to assess off target effects of Mtz. Fish in Mtz groups were given 7mM of Mtz dissolved in DMSO. Fish from each group began treatment starting from SSL 7.0 and continued to receive treatment until reaching SSL 12.0. During treatment fish were fed 3 times per day using 1-2 drops of brine shrimp. Upon reaching SSL 12.0, fish were euthanized using tricaine and embedded into a 1% agarose gel in a sterile petri dish. Colored images were taken using a Koolertron LCD digital microscope.

Overnight Treatments: Metaphocytes were ablated overnight for time lapse imaging using fish injected with aox5:palmEGFP into *Tg(grn2:EGFP-v2a-nfsB)*. Fish were treated with either 10 mM Mtz or DMSO as a control. MMP9+ cells were ablated for time lapse imaging using fish injected with aox5:palmEGFP into *Tg(mmp9:EGFP-v2a-nfsB)*. Fish were treated with either 5 mM Mtz or DMSO as a control. MMP9 inhibition for overnight time lapse was using fish injected with aox5:palmEGFP into *Tg(mpeg1:tdtomato)* or WT. Fish were treated with either 0.1 mM MMP9 inhibitor II or DMSO as a control.

### Time-lapse and Still Imaging

*Ex vivo* imaging of pigment cells and macrophages was done using a Leica TCS SP8 confocal microscope equipped with a resonant scanner and two HyD detectors. Time-lapse images taken at 5-minute intervals over a span of 10 hr. Overnight time-lapse imaging was performed when larvae reached SSL 7.5 except where indicated (Eom et al., 2015; Eom and Parichy, 2017). NTR ablation experiments were performed by administering either 10 mM Mtz into *Tg(grn2:EGFP-v2a-nfsB)* or 5 mM, *Tg(mmp9:EGFP-v2a-nfsB)* overnight for airineme extension frequency measures.

### Cell Counts and Distributions

Pigment Pattern: Fish were imaged upon reaching SSL 12.0. Images were taken of the entire trunk and were later cropped to include only the area underneath the dorsal fin. Total melanophore counts and melanophores in the interstripe region were quantified using the cell counter plugin from ImageJ. Total number of melanophores and melanophores in the interstripe region were averaged.

Developmental Stage Cell Population Counts: Dendritic and amoeboid cell populations were quantified using the cell counter plugin from ImageJ. Dendritic cells were determined by several characteristics including *mpeg1+* (amoeboid & dendritic macrophages) reporter expression level, cell morphology, migration speed, branching. Total numbers for each group were averaged across individuals. *gnr2+* cells (metaphocytes) were also quantified using the cell counter plugin of ImageJ and averaged across individuals.

Axial Localization of skin-resident macrophages: Localization of amoeboid and dendritic populations was determined using z-stack information provided by images taken on Leica TCS SP8 confocal microscope. Distance from the hypodermis was measured and determined by co-injected aox5:palmEGFP+ cells. Thus, smaller numbers indicate closer to the hypodermis and larger numbers closer to the epidermis (Fig. 6).

### Statistical analysis

Statistical analyses were performed using GraphPad Prism software version 9.0 for Mac (GraphPad Software, San Diego, CA, USA). Frequency data for behaviors of individual cells or projections were assessed by single or multiple factor maximum likelihood. Continuous data were displayed as mean ± SEM, using an unpaired two-tailed Student’s t-test or analyses of variance.

| Tg Line/Construct Used | Name | Lab Origin |
| --- | --- | --- |
| *TgBAC(grn2:EGFP-NTR)* | hkz034 | Wen Lab |
| *Tg(grn2:EGFP;mpeg1:LRLG)* | hkz015<br>(for *mpeg1:LRLG*) | Wen Lab |
| *Tg(cldnh:EYFP-v2a-nfsB)* | **This paper** | cldnh:eGFP construct from Wen lab was subcloned to generate cldnh:EYFP-v2a-nfsB and screened for Tg |
| cldnh:eGFP (construct) | Construct:<br>*Tg(cldn3c:GFP)* | Wen Lab |
|  | Line: hkz021 |  |
| *TgBAC(mmp9:eGFP)* | tyt206 | Kawakami Lab |
| *TgBAC(mmp9:eGFP-nfsB)* | tyt207 | Kawakami Lab |
| *Tg(mpeg1:Brainbow)* | W201 | Ramakrishnan Lab |
| *TgBAC(tyrp1b:palmmcherry)* | wprt11 | Parichy Lab |
| *mpeg1:NTR-EYFP* | w202 | Moon Lab |
| *Tg(TNFa:emGFP)* | **This paper** | TNFa:emGFP construct was gifted from Roh-Johnson lab and we generated a Tg line with injections. |
| *cldnh:mcherry* (construct) | **This paper** | *cldnh:eGFP* construct from Wen lab was subcloned to generate *cldnh:mCherry* |
| *aox5:palmEGFP* (construct) |  | Parichy Lab |

## Acknowledgements

We thank Dr. T. Schilling for critical reading and editorial input and Dr. J. Lee for valuable discussion. We acknowledge support from NIH MIRA R35GM142791 to DSE. We thank the Dr. Zilong Wen in Hong Kong University of Science and Technology for sending us *Tg(grn2:EGFP), Tg(grn2:EGFP-nfsB), cldnh:EGFP*, and National BioResource Project, Zebrafish, Core Institution (NZC) in Japan for providing us *Tg(mmp9:EGFP)* and *TG(mmp9:EGFP-nfsB)* lines. We also thank Dr. Minna Roh-Johnson at the University of Utah for sending us *TNFa:emGFP* construct.

## Author contributions

RLB and DW performed experiments and analyzed data. DSE designed the experiments, performed experiments, analyzed data, and wrote the manuscript.

### Ethics

All animal work in this study was conducted with the approval of the University of California Irvine Institutional Animal Care and Use Committee (Protocol #AUP-22-023) in accordance with institutional and federal guidelines for the ethical use of animals.

**Supplementary Figure 1.**
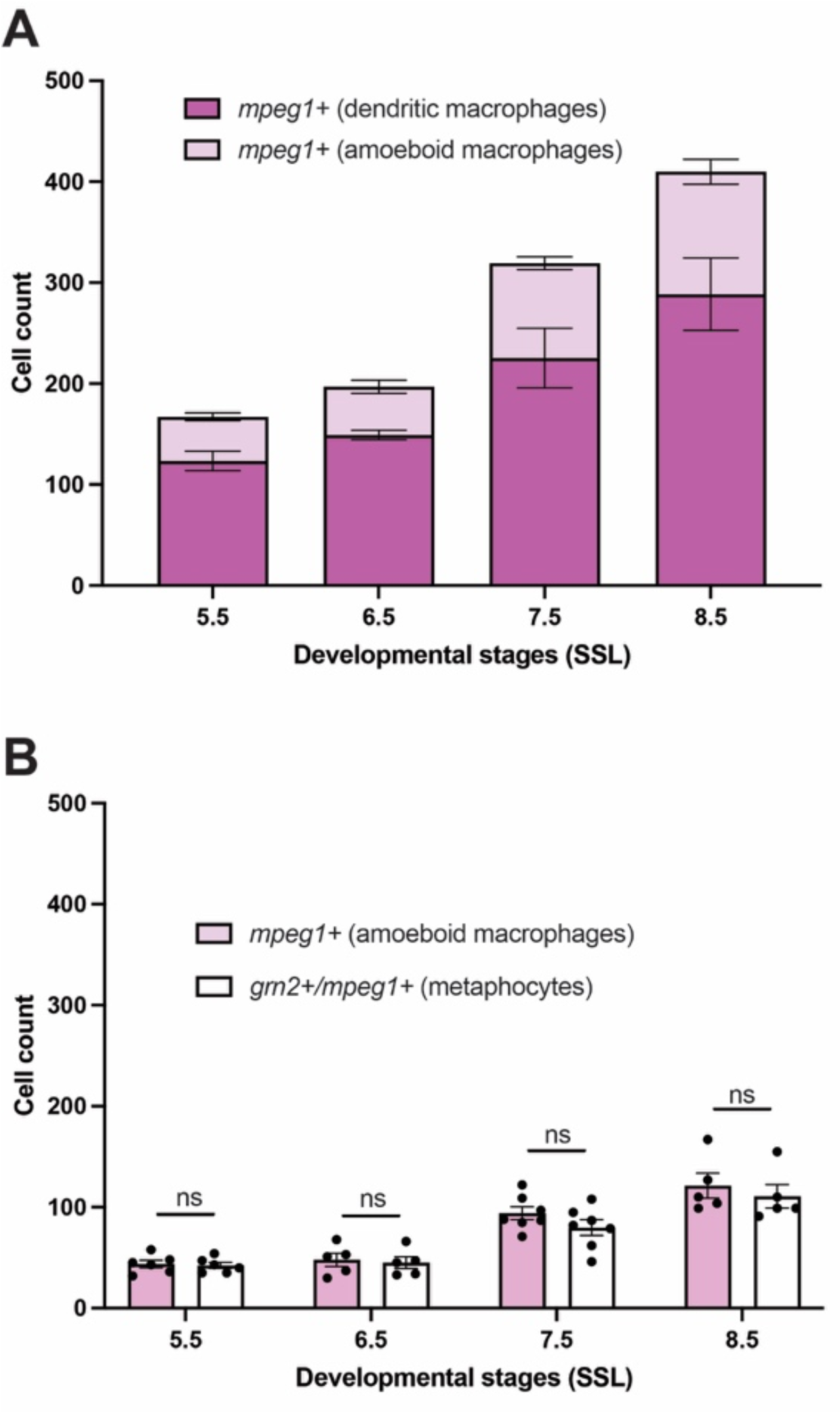
Skin-resident macrophage populations in various developmental stages in zebrafish. (A) Developmental stage-dependent cell counts in both dendritic and amoeboid macrophage subpopulation. The proportion of amoeboid macrophages gradually increases as fish develops (∼24% at SSL5.5 to ∼29% at SSL8.5), but no dramatic expansion at SSL 7.5 (n=8 trunks at SSL5.5, 5 at SSL 6.5, 7 at SSL 7.5, 6 at SSL8.5). (B) Numbers of amoeboid macrophages (*mpeg1+)* and metaphocytes (*mpeg1+/grn2+)* are not differ in various developmental stages (n=5 trunks at SSL 5.5, 5 at SSL 6.5, 7 at SSL 7.5, 5 at SSL 8.5).

**Supplementary Figure 2.**
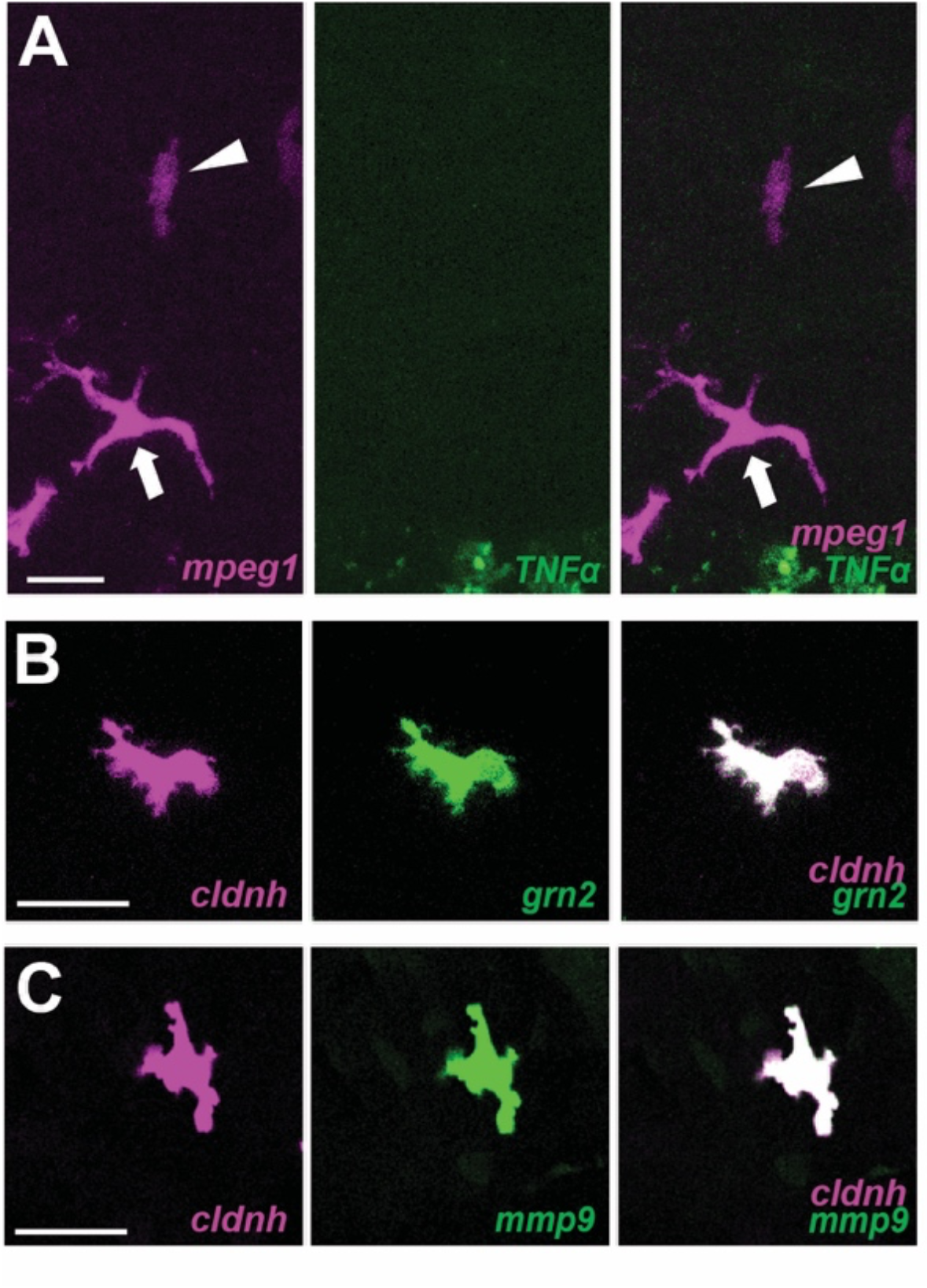
Amoeboid macrophage subpopulation. (A) Neither amoeboid (arrowhead) and dendritic (arrow) macrophages are *TNFα* positive. (B) Amoeboid macrophage subpopulation express both metaphocyte markers (*grn2* and *cldnh*). (C) Amoeboid macrophages express metaphocyte marker, *cldnh* and *mmp9*.

**Supplementary Figure 3.**
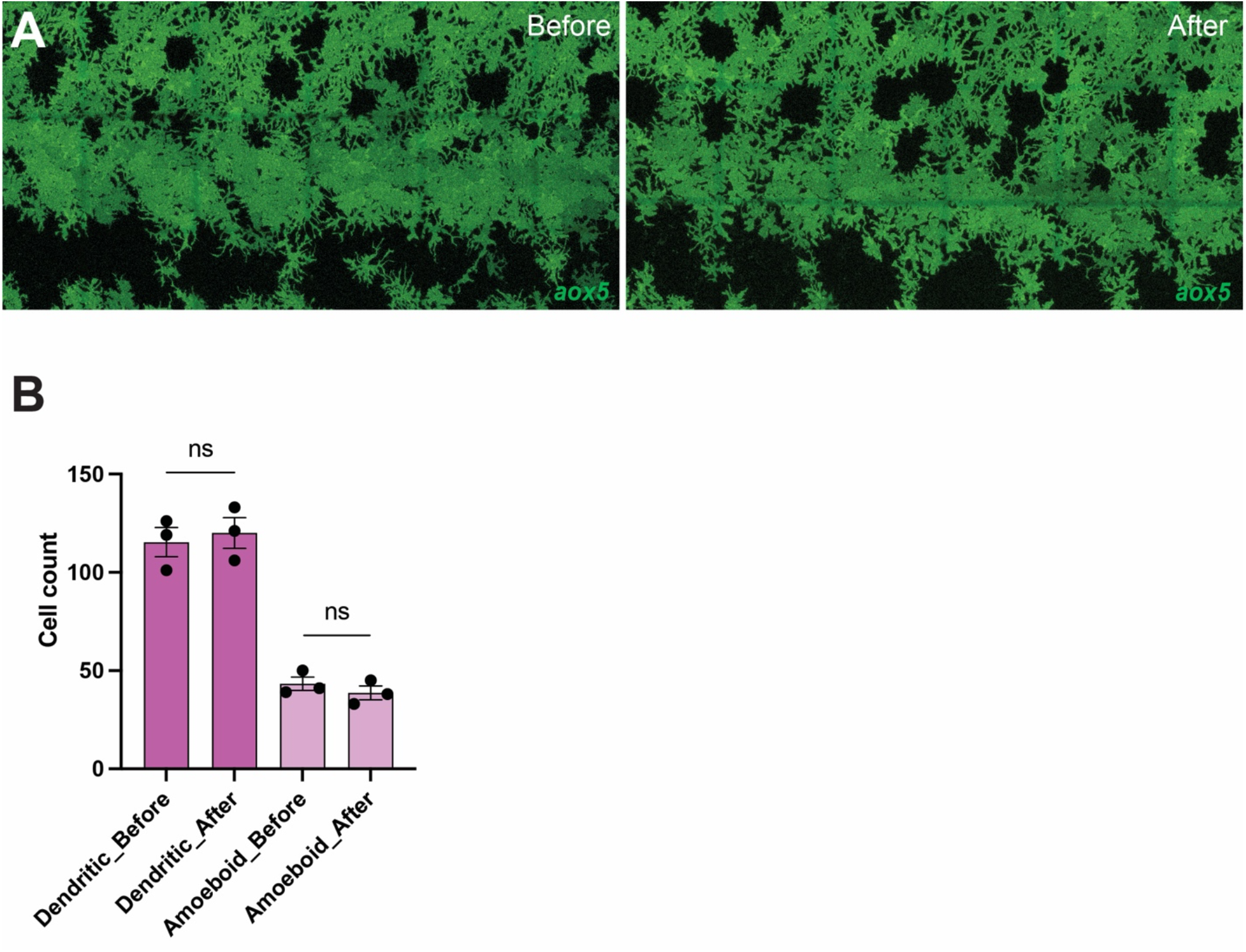
MMP9 inhibitor treatment did not affect the number of xanthophore-lineage cells and *mpeg1+* macrophages. (A) Xanthophore-lineage cells are unaffected by MMP9 inhibitor treatment. (B) Neither dendritic nor amoeboid macrophage numbers were changed upon MMP9 inhibitor treatment.

**Video 1. Skin-resident macrophages in developing zebrafish**

Arrowheads indicate amoeboid macrophages and arrows for dendritic macrophages. Note that amoeboid macrophages migrate faster than dendritic. Macrophages were labelled with *Tg(mpeg1:tdtomato)*, pseudo-colored in magenta. Time stacks indicated at the right top corner. 5 min interval for 1 hr 20 mins.

**Video 2. Airineme pulling macrophage subpopulation**

Airineme vesicles pulled by amoeboid macrophage subpopulation (arrowhead). Xanthophore-lineage cells (green) were labelled with aox5:palmEGFP injection into *Tg(mpeg1:tdtomato)*, pseudo-colored in magenta. Time stacks indicated at the right top corner, 5 min interval for 30 mins.

**Video 3. Airineme extension by metaphocytes**

Airinemes pulled by metaphocytes (arrowhead, *grn2+*). Xanthoblasts were labelled with aox5:palmEGFP injection into *Tg(grn2:EGFP)*. Metaphocytes were distinguishable by their morphology and fast migration behavior. Time stacks indicated at the right top corner, 5 min interval for 25 mins. Scale bar: 10μm.

**Video 4. Metaphocyte or mmp9+ cell ablation reveals the requirement for airineme extension**

Left, airinemes were extended frequently (arrowheads) in the DMSO control; Middle, no airinemes were observed in the metaphocyte ablated embryo, Right, no airinemes were observed in the mmp9+ cell ablated embryo. Xanthoblasts (green) were labeled with F0 injection of *aox5:palmEGFP* construct into *Tg(grn2:EGFP-v2a-nfsB)* or *Tg(mmp9:EGFP-v2a-nfsB)*. Time stacks are indicated at the top right corner of the control movie, 5 min intervals for 5 hrs.

**Video 5. MMP9 inhibitor treatment inhibits airineme extension**

Left, airinemes were extended frequently (arrowheads) in the DMSO control; Right, no airinemes were observed in the MMP9 inhibitor treated embryo, Xanthoblasts (green) were labelled with F0 injection of *aox5:palmEGFP* construct into WT(ABb). Time stacks are indicated at the top right corner of the control movie, 5 min intervals for 5 hrs.

